# Enhancing network activation in Natural Killer cells: Predictions from *in silico* modeling

**DOI:** 10.1101/395756

**Authors:** Sahak Z. Makaryan, Stacey D. Finley

## Abstract

Natural killer (NK) cells are part of the innate immune system and are capable of killing diseased cells. As a result, NK cells are being used for adoptive cell therapies for cancer patients. The activation of NK cell stimulatory receptors leads to a cascade of intracellular phosphorylation reactions, which activates key signaling species that facilitate the secretion of cytolytic molecules required for cell killing. Strategies that maximize the activation of such intracellular species can increase the likelihood of NK cell activation upon contact with a cancer cell, and thereby improve efficacy of NK cell-based therapies. However, due to the complexity of intracellular signaling, it is difficult to deduce *a priori* which strategies can enhance species activation. Therefore, we constructed a mechanistic model of the CD16, 2B4 and NKG2D signaling pathways in NK cells to simulate strategies that enhance signaling. The model predictions were fit to published data and validated with a separate dataset. Model simulations demonstrate strong network activation when the CD16 pathway is stimulated. The magnitude of species activation is most sensitive to the receptor concentration and the rate at which the receptor is deactivated. Co-stimulation of CD16 and NKG2D *in silico* required fewer ligands to achieve half-maximal activation than other combinations, suggesting co-stimulating these pathways is most effective in activating the species. We applied the model to predict the effects of perturbing the signaling network and found two strategies that can potently enhance network activation. When the availability of ligands is low, it is more influential to engineer NK cell receptors that are resistant to proteolytic cleavage. In contrast, for high ligand concentrations, inhibiting phosphatase activity leads to more activation. The work presented here establishes a framework for understanding the complex, nonlinear aspects of NK cell signaling and provides detailed strategies for enhancing NK cell activation.

## 1 INTRODUCTION

Natural killer (NK) cells are immune cells that can eliminate cancer cells upon cell contact (1–3). NK cells express a repertoire of stimulatory receptors that mediate the release of cytotoxic chemicals when stimulated by antibodies or by cells that express stimulatory ligands. The activation of such receptors induces intracellular signaling through a cascade of phosphorylation reactions, which ultimately leads to NK cell activation, degranulation and cancer cell death. This innate ability for cancer cell elimination has spurred an interest in research (3–5) to better understand NK cell activation. It is believed that enhancing NK cell activation could proportionally enhance cancer cell killing and thereby improve patient outcomes in the clinic. Given that cancer cell killing is initiated via activation of NK cell stimulatory receptors, it is important to understand how the signal propagates and activates the downstream species that contribute to NK cell activation. Therefore, researchers (1–4,6,7) have studied NK cell signaling and reported which species are activated downstream of the stimulatory receptors. Such findings are crucial in understanding how NK cell activation proceeds on the molecular level.

However, due to the natural complexity and nonlinearity underpinning intracellular signaling, it is difficult to deduce how NK cell signaling can be modulated to enhance cell activation. Mathematical models are valuable in these contexts in that they enable us to untangle such complicated system behavior and predict the system’s response to a wide variety of perturbations (8–14). For example, work by Das demonstrated how receptor-ligand interactions impact NK cell activation and the various NK cell responses induced by strong and weak stimulatory ligands (9). Mesecke and colleagues showed that the physical association of Src family kinases (SFK) with a stimulatory receptor is essential for NK cells to promote a cytotoxic response, and that the activation of the signaling species Vav correlates with NK cell cytotoxicity (10). Nevertheless, the question of which strategies enhance NK cell signaling (and why) remains open. Additionally, the previous models did not determine which molecular perturbations or which pathways should be co-stimulated to optimally activate the NK stimulatory network.

Here, we developed a molecularly detailed, experimentally validated mechanistic model of NK cell signaling to address the above questions. The CD16, 2B4 and NKG2D stimulatory pathways were modeled in this study as these pathways contribute to NK cell activation in different ways (4,15,16). CD16 is an Fc receptor that binds to the constant region of antibodies. This implicates CD16’s activation in antibody-dependent cell-mediated cytotoxicity (ADCC). Its cytoplasmic domain is associated with CD3ζ, which contains immunoreceptor tyrosine-based activation motifs (ITAM). 2B4 is part of the signaling lymphocytic activation molecule (SLAM) family of receptors, and its cytoplasmic tail contains four immunoreceptor tyrosine-based switch motifs (ITSM). The ligand for 2B4, CD48, is expressed by cells of hematopoietic origin. This suggests 2B4 may play a role in regulating hematopoietic processes. NKG2D belongs to the family of C-type lectin-like receptors. It associates with the adaptor protein DAP10, which has an activation motif that is similar to the CD28 T cell co-receptor. NKG2D binds to ligands typically expressed by cells that have undergone transformation, which implicates this receptor in the elimination of tumors.

Ligand binding to the CD16, 2B4 and NKG2D receptors initiates intracellular signaling. The PI3K-Akt, SLP76-Vav-Erk and PLC*γ* networks are all activated upon CD16, 2B4 and NKG2D stimulation (17). In NK cell biology, the PI3K-Akt pathway promotes cell survival, while Erk activation is correlated with cell proliferation. SLP76 and Vav activation are necessary for actin remodeling and the formation of the immunological synapse. Lastly, PLC*γ* activation induces the release of intracellular calcium ions, which subsequently contributes to cell activation. The combination of these intra-cellular reactions is necessary to activate NK cells. Many of the downstream reactions are common between the pathways with only subtle differences. For example, 2B4 does not induce Akt phosphorylation (6,7). Additionally, 2B4 and NKG2D specifically lead to phosphorylation of the Y113 and Y128 sites on SLP76, respectively, while CD16 induces phosphorylation of both sites (6). Also, CD16 induces ZAP70 and LAT activation, while 2B4 and NKG2D do not. Thus, these pathways are interconnected, and understanding the dynamics of the concentrations of the molecular species involved in the signaling pathways requires in-depth analyses.

In the present study, we use mathematical modeling to characterize and compare the signaling dynamics of the CD16, 2B4, and NKG2D pathways with respect to their magnitude of activation of the network. Furthermore, we identify which signaling species and parameters influence the magnitude of network activation and which combinations of receptor co-stimulation most potently activate the network. *In silico* perturbations of the stimulatory network demonstrate the strategies that effectively increase network activation, including which signaling species to target and how to modify the species. In total, the model predictions can be used for engineering NK cells with enhanced signaling, which is needed for cell activation and ultimately target cell killing.

## 2 METHODS

### Model construction

We constructed an ordinary differential equation (ODE) model to predict the concentrations of the molecular species in the CD16, 2B4, and NKG2D pathways in NK cells (**Figure 1**). The model is provided in Supplementary File S1 and the list of model species, reactions and parameters are provided in Supplementary File S2. The rates of the biochemical signaling reactions were represented using Michaelis-Menten reactions. Arriving at the current model structure was an iterative process where we fitted several model types (e.g., Michaelis-Menten kinetics vs. Mass Action kinetics, including vs. excluding the phosphatases, including vs. excluding non-specific decay rate) to the experimental data and selected the model structure that generated the lowest error. We constructed the model using BioNetGen (18) and simulated it in MATLAB (MathWorks).

**Fig. 1.**
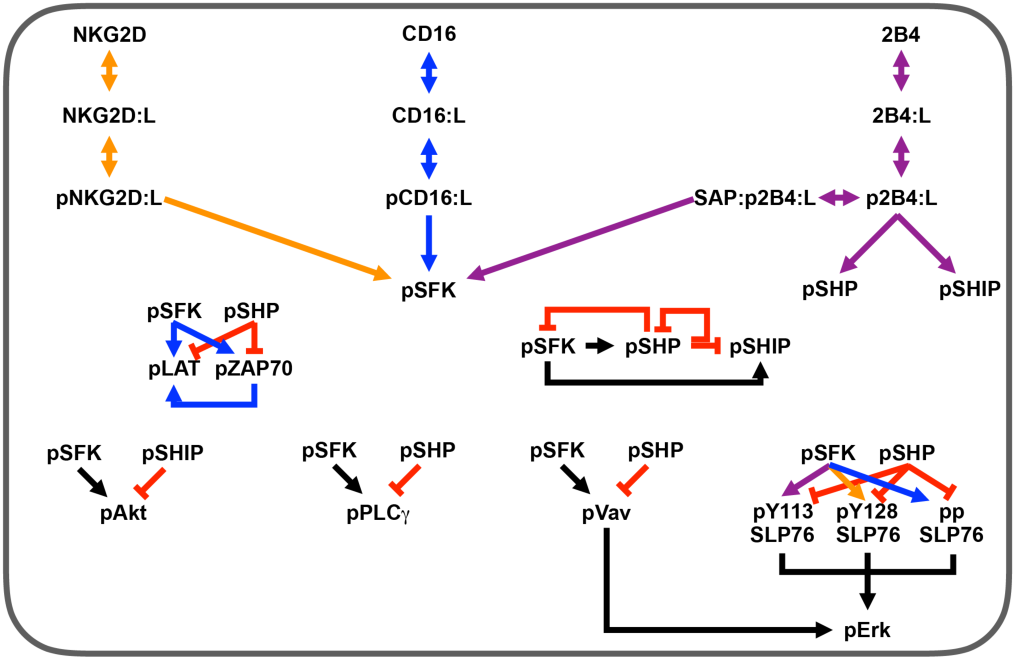
Model schematic. Reaction network for three stimulatory receptors expressed on the surface of NK cells: CD16, 2B4 and NKG2D. These receptors promote signaling species that mediate NK cell activation: SFK, Erk, Akt and PLC*γ*. Arrows indicate stimulation, while red crossbars indicate inhibition. Orange arrows are specific to the NKG2D pathway; blue, CD16 pathway; purple, 2B4 pathway; black, all pathways.

The final model contains 83 parameters and 36 species, including the three NK cell receptors. Each receptor binds to its ligand and forms a receptor-ligand complex that allows the receptor to become phosphorylated by basally active Src family kinases (SFK). Then, the ligand-bound phosphorylated receptor serves as the catalyst for converting SFK from a basally the active state to a fully active state (pSFK). Fully active SFK mediates the phosphorylation (activation) of a number of downstream signaling species, including LAT, ZAP70, PLC*γ*, Vav, SLP76, Akt, and the phosphatases SHP and SHIP. Moreover, the stimulation of 2B4 can lead to activation of the phosphatases independent of pSFK (17). Phosphorylated ZAP70 promotes activation of LAT. The inhibitory species, phosphatases SHP and SHIP, provide negative feedback to prevent overactivation (19). The catalysts for Erk phosphorylation are the phosphorylated forms of SLP76 and Vav. These species are upstream inputs to the MAPK pathway (9,10).

The initial concentrations of the species in our system were extracted from the literature (20–23). We simulate the dynamics of the signaling network for 60 minutes, to focus on the initial stimulus. Given this time scale, we assume that the synthesis of species is negligible compared to the rates of the phosphorylation and dephosphorylation reactions (9,10). Lastly, we included a non-specific degradation reaction of the phosphorylated species in the system to account for degradation, dilution and disappearance of the active species (6,24–27).

### Data collection and processing

We trained and validated the mathematical model using experimental data extracted from the literature. The raw data and our data processing procedure are provided in Supplemental File S3. To control for variations in the experimental conditions, we only used data from published studies where (1) the antibodies used for CD16, 2B4 and NKG2D stimulation were of the same concentration (10 *μ*g/mL) and from the same vendor, and (2) the cell types used in these studies were primary NK cells. Immunoblot images from these published studies were analyzed and processed using ImageJ (28). Specifically, ImageJ provides a measure of the optical density for any pre-defined rectangular space of an image in grayscale, where the estimated optical densities range from 0 – 225 (black to white, respectively). Protein bands in Western blots were analyzed to estimate their optical density. To control for immunoblot variations across the experiments, we subtracted the optical density measurement of the Western blot gel background from the optical density measurements of all protein bands in the same gel. Furthermore, for a single protein, the optical density measurement of the zeroth time point was also subtracted from the optical density measurements of the remaining time points. This procedure, which follows the documented ImageJ usage protocol, standardizes the experiments for comparison and controls for the background and zeroth time point measurements. In total, the model was trained to 64 data points. Additionally, the model was validated against 32 data points. The signal intensity 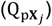 of a given phosphorylated species (p**X**) at the *j*^*th*^ time point is calculated as:

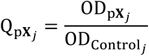

where 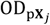 and 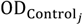 are the optical density values of the phosphorylated species and a loading control, respectively, at the *j*^*th*^ time point. Furthermore, the signal intensity (Q_p**X**_) was normalized to a single (reference) time point 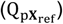 by calculating the percent change (%Δ_p**X**_) as follows:

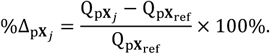

### Parameter estimation

The parameters were estimated using a Bayesian perspective (29), where we maximized the posterior density (*f*(**θ**|**Data**)) of the parameters (**θ**) given the data (**Data**) via the Metropolis-Hastings algorithm (29,30). Briefly, as the parameters in the present model are known to follow a lognormal distribution (29,30), we used this distribution as our prior (*f*(**θ**)). Moreover, to simplify estimation, we assume parameters of the same kind (e.g., *k*_cat_ parameters) are independent and drawn from an identical distribution (IID). We further assume the error between the model predictions and the data (i.e., the likelihood function *f*(**Data**|**θ**)) follows a Gaussian distribution centered at zero with some variance (*σ*^2^). Since the proposal distribution from which the parameters are sampled from must have proper support, we again utilized the lognormal distribution as it has domain over the positive real line. We ran this fitting algorithm 200 times using randomized initial guesses, with each independent run simulated for 10,000 iterations. We discarded the first 9,000 iterations, where the algorithm searches the parameter space to maximize the posterior density. The last 1,000 iterations are where we begin to sample from the posterior distribution, and we used those 1,000 iterations to identify the best fit parameter sets (i.e., those with the lowest error between model predictions and experimental data). The best fit parameter sets were used for model simulations.

### Construction of magnitude of network activation metric

We defined network activation to allow us to compare the magnitude of signaling across the three pathways (CD16, 2B4 and NKG2D). While the individual phospho-species are known to contribute to specific cellular functions involved in NK cell activation (6,17,26,31–34), the scope of the current work is to compare the effectiveness of stimulating one pathway versus another. Thus, we first determine which species are important to consider in terms of activation of the network.

Based on literature evidence, we determined the following five species, which are common to all three pathways, are crucial for activating the NK cell based on experimental studies: (1) pErk, (2) pAkt, (3) pPLC*γ*, (4) pVav and (5) pSLP76. The magnitude of network activation must relate to the magnitude of activation of the above species. Hence, we concatenate the above species’ concentrations over time into a vector,

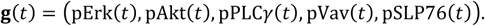

Since the phospho-species’ concentrations are continuous with respect to time *t*, **g**(*t*) is also continuous with respect to *t* and thus measurable (35). By construction, the arc **g** is a function that maps the time interval (in minutes) [0,60] in a subset of ℝ^5^; more specifically, **g**: [0,60] → ℝ^5^. We used the Bochner-norm to define the magnitude of **g**(*t*) (i.e., ‖**g**(*t*)‖). Here, the Bochner-norm of **g**(*t*) is defined by:

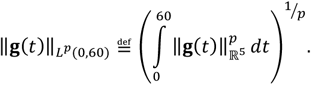

Since all norms in ℝ^n^ form equivalence classes (36), we set *p* = 1 and use the *L*^1^norm on ℝ^5^ for simplicity. In addition, each component *g*_*i*_(*t*) is non-negative and has finite measure, and the sum of the components, for all *t* ∈ [0,60], is finite. Therefore, Fubini’s theorem applies (35) and we can switch the order of summation and integration. Finally, the Bochner-norm (in *L*^1^) of **g**(*t*) defined here is

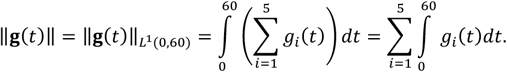

Thus, we arrive at the above metric for the magnitude of network activation, which is simply the sum of the magnitude of activation of the individual phospho-species, as given by the area under the curve for the species’ concentration profile. We used the MATLAB function *trapz* (which uses trapezoidal numerical integration) to estimate the area under the curve for each component of *g*(*t*).

### Clustering and principal component analysis

We used the built-in MATLAB functions *kmeans* and *pca* to perform *k*-means clustering and principal component analysis, respectively. Briefly, *k*-means clustering (37) allows us to partition a given dataset into *k* clusters using the (default) Euclidean distance metric. Principal component analysis (38) enables us to project a given dataset on to a new coordinate system where each coordinate is a linear combination of the original variables in the dataset. Moreover, the principal components (i.e., new coordinates) are selected such that they maximize the total variance in the data. These approaches are used to determine which estimated parameter sets are similar to one another.

## 3 RESULTS AND DISCUSSION

### 3.1 Model of NK cell signaling matches experimental data

We generated a mathematical model of NK cell signaling that includes three main pathways: CD16, 2B4 and NKG2D. When these receptors are stimulated, they activate the cell via cascades of phosphorylation reactions (**Figure 1**): activation of the Src family kinases (SFK), facilitated by the ligand-bound phosphorylated receptors, catalyzes the activation of the Akt, SLP76-Vav-Erk, and PLC*γ* pathways. We simulated these reactions in the form of nonlinear ordinary differential equations (ODEs) using established Michaelis-Menten kinetics. The model is provided in Supplementary File S1. The model was calibrated to immunoblot data (6,24,25,27), where we quantified the temporal change in the optical density of protein bands from images of immunoblot experiments using ImageJ (28). Specifically, we used the normalized levels of the following phosphorylated species: pSFK, pZAP70, pLAT, ppSLP76, pPLC*γ*, pVav, pErk, pAkt, and SLP76 phosphorylated at Y113 and Y128. We calibrated the model predictions by estimating the parameter values using a Bayesian perspective (29), and by implementing the Metropolis-Hastings algorithm (see Methods). In brief, the model parameters (83 in total) were estimated 200 times using randomized initial guesses by fitting to experimental data. Moreover, the model predictions were validated using a separate dataset. The combined error for each run can be found in **Figure S1**. We proceeded with the 14 best parameter sets that provided the lowest total error and simulated the model using these sets.

Interestingly, in initial simulations, we found that the 14 parameter sets led to different responses with respect to network activation. To determine the dominant behavior generated by the model, we clustered the network activation predicted by the 14 sets using the *kmeans* and *pca* functions in MATLAB (see Methods). Our results are shown in **Figure S2**, where we identified three unique clusters that correspond to the degree of species activation (i.e., response) predicted by the model. To ensure that the model predictions agree with experimental observations, we discarded the parameter sets that yielded predictions inconsistent with NK cell signaling and cytotoxicity studies (6,24–27,33,39,40). Specifically, we removed parameter sets that induced a low amount of species activation (i.e., < 1% of the species’ initial concentration was activated) and that did not show a dosedependent response when the ligand concentrations were changed. This refined the 14 parameter sets down to five, which are found in the medium response cluster in **Figure S2**. The parameter distributions for the best set (i.e., lowest total error) of the five are shown in **Figure S3** using the final 1,000 iterations, illustrating that the parameters are well behaved: the distributions are unimodal, and the values lie within a tight range. We used the last 1,000 iterations from parameter estimation to simulate the model.

The simulated concentration profiles are consistent with the training data (**Figure 2A – H**). These results demonstrate the model predictions are in accord with the experimental data for mono-stimulation of CD16 (blue lines), 2B4 (purple lines) and NKG2D (orange lines). This is expected, since those data were used in model training to determine the parameter values.

**Fig. 2.**
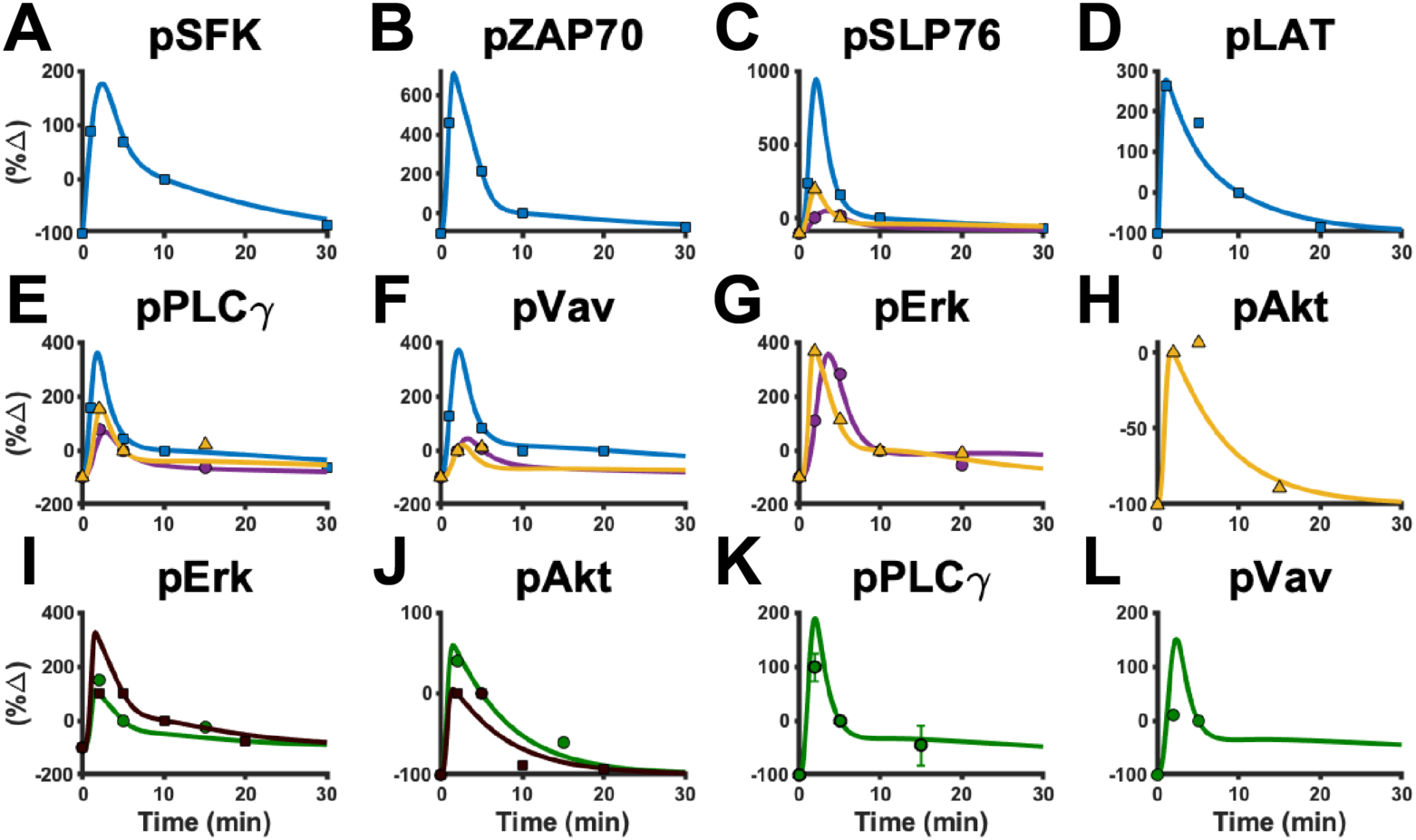
Model calibration and validation. The model was fit to experimental data for (**A**) pSFK, (**B**) pZAP70, (**C**) pSLP76, (**D**) pLAT, (**E**) pPLC*γ*, (**F**) pVav, (**G**) pErk and (**H**) pAkt. The model predictions were validated against separate data for (**I**) pErk, (**J**) pAkt, (**K**) pPLC*γ* and (**L**) pVav under co-stimulation of 2B4 and NKG2D. Blue: CD16 pathway. Purple: 2B4 pathway. Orange: NKG2D pathway. Green and Brown: 2B4 and NKG2D co-stimulation from separate experiments. Note that the green and brown lines represent independent Western blot experiments that only differ in the time-points of data collection. Solid lines: mean model predictions from 1,000 parameter estimates. Shaded area: standard deviation of mean model predictions. Squares, circles and triangles: experimental data.

To validate the model, we compared the model predictions to separate experimental data not used during training. In particular, we quantified the optical density of intracellular species from immunoblot images when 2B4 and NKG2D were simultaneously stimulated at equal ligand concentrations (24,26,33,39,40). The results from model validation are shown in **Figure 2I – L**. The model captures the signaling dynamics of several species upon co-stimulation of 2B4 and NKG2D. Altogether, this validated model allows us to perform simulations and make meaningful comparisons amongst the pathways.

### 3.2 Baseline network activation is greatest when the CD16 pathway is stimulated

In addition to the amount of activation of the phosphorylated species, we were interested in quantifying the magnitude of activation of the network induced by each pathway. Here, we use the norm of the vector-valued function **g**(*t*), where each component of this vector is the time evolution of the concentration of the five species considered to be necessary for NK cell activation (pErk, pAkt, pPLC*γ*, pVav and pSLP76 (see Methods for derivation)). The model was simulated for 60 minutes using 6.67 × 10^−2^ *μ*M of ligand, the same concentration used in the experimental studies to train and validate the model (6,24–27,33,39,40). We used the last 1,000 iterations from the best set obtained from parameter estimation.

The magnitude of network activation is greatest for the CD16 pathway compared to stimulation of 2B4 and NKG2D at equal ligand concentrations (**Figure 3**). Interestingly, each pathway activates the network differently (**Figure S4**). For example, CD16 induces more activation of pSLP76 (**Figure S4A**), whereas stimulation of the NKG2D pathway activates pVav (**Figure S4B**) and pPLC*γ* (**Figure S4C**) to a greater extent. In contrast, the pathways show no significant difference with respect to pErk (**Figure S4D**) and pAkt (**Figure S4E**) activation. These results support our systems-level evaluation of the network, as focusing on a single species does not fully represent the effects of stimulating an NK receptor. The baseline model is useful in allowing us to quantitatively interpret the results from the experimental studies used to train the model, where the signaling species concentrations were not measured directly. Overall, the baseline model predicts CD16 stimulation leads to a greater activation of the stimulatory network and that the magnitude of activation of the phosphorylated species varies depending on the pathway being stimulated.

**Fig. 3.**
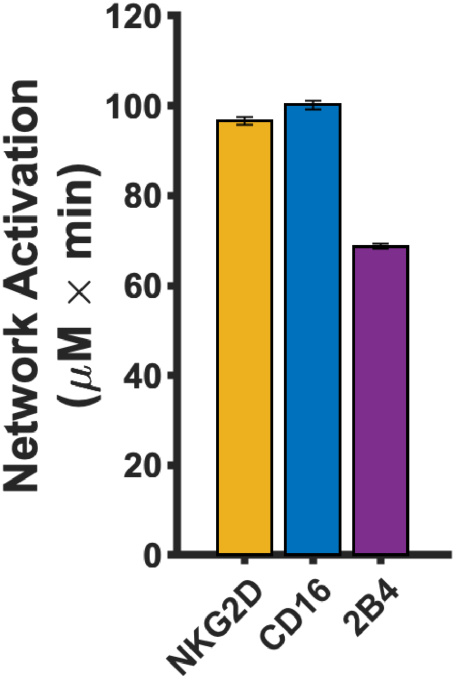
Baseline network activation of individual receptors. The magnitude of network activation induced by mono-stimulation of NKG2D (orange), CD16 (blue) and 2B4 (purple). Bars represent the mean model prediction from the 1,000 parameter estimates and the error bars represent one standard deviation.

### 3.3 Receptor characteristics significantly influence the network activation

The stimulation of NK cell receptors with ligands (model input) leads to activation of the signaling species (model output). Ultimately, we wish to understand the output of the system as a function of its input. To achieve this, we varied the ligand concentration *in silico* from 6.67 × 10^−5^ *μ*M up to 66.7 *μ*M and simulated the model for 60 minutes to observe how the magnitude of network activation changes. Such a wide range is typically implemented in experimental studies as it allows researchers to understand how a system behaves as its input changes in magnitude (12). As such, our results in **Figure 4** show how the predicted magnitude of network activation changes as ligand concentrations change. We present the mean model prediction (solid line) using the final 1,000 iterations from the best parameter set, along with its standard deviation (shaded area).

**Fig. 4.**
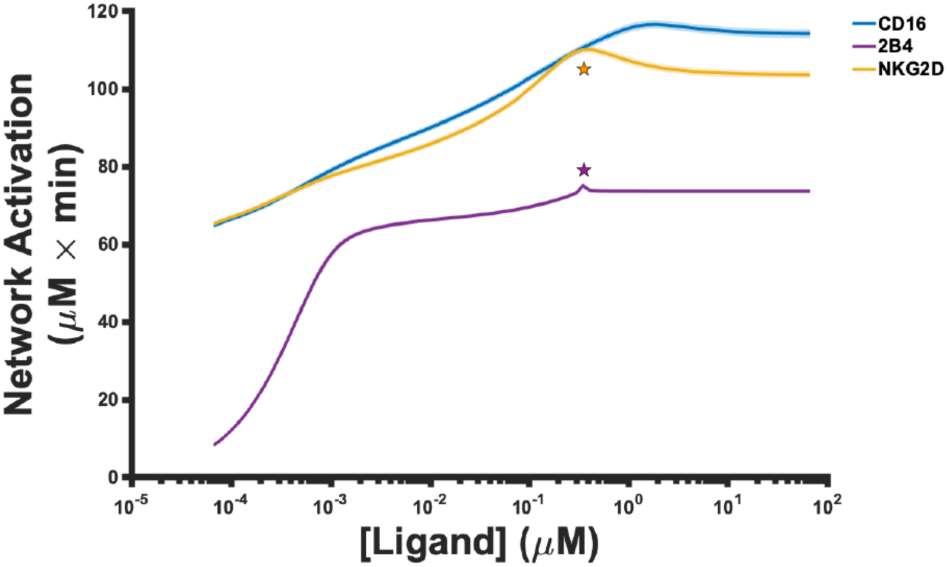
Network activation as a function of ligand stimulation. The mean value (solid line) of the magnitude of network activation from the 1,000 parameter estimates, along with one standard deviation (shaded area), is shown for stimulation of CD16 (blue), 2B4 (purple) and NKG2D (orange).

The model predicts that, in general, the magnitude of network activation increases as more ligands are introduced into the system (**Figure 4**). For all ligand concentrations we simulated, the model predicts that the magnitude of network activation is always greater when either the CD16 or NKG2D pathways are stimulated, compared to the stimulation of 2B4. Our results suggest that mono-stimulation of 2B4 induces weak activation of the stimulatory network.

Interestingly, we observed an unexpected sharp peak in network activation (**Figure 4**; purple star) at approximately 0.3 *μ*M of ligand during stimulation of 2B4. Given the detailed nature of the model, we could apply it to investigate the cause of the peak. The ligand concentration at which the peak occurs is numerically close in value to the concentration of 2B4 receptor in the model (0.353 *μ*M). Thus, we simulated the model using different concentrations of 2B4 to determine if the peak in network activation is due the receptor’s concentration (**Figure S5A**). Indeed, the concentration of 2B4 sets the threshold for network activation since we observe more network activation when we increase the receptor’s concentration accordingly. In addition, network activation induced by 2B4 is maximal when the ligand concentration reaches the same level as the receptor.

Similar to 2B4-mediated network activation, the model predicts a peak in NKG2D-mediated network activation (**Figure 4**; orange star). Again, the peak occurs when the ligand concentration is approximately the same as the concentration of NKG2D in the model (0.303 *μ*M). We simulated the model using different concentrations of NKG2D as we did above (**Figure S5B**). We observed that the concentration of NKG2D sets the upper bound on network activation, where the peaks occur when the concentration of ligand is near the concentration of the receptor.

In the case of NKG2D, unlike 2B4, there is a notable decrease in network activation once the ligand concentration is greater than the concentration of the receptor (**Figure 4**; orange star). To better understand this observation, we varied the parameters regulating NKG2D phosphorylation and dephosphorylation. The model predicts that the catalytic rate constant for phospho-NKG2D dephosphorylation (*k*_*cat*_*_pNKG2D_pSHP* in the model) is responsible for this behavior. Namely, once we increased the value of this parameter by 10-fold (**Figure S5C**) or 100-fold (**Figure S5D**) from its baseline value (14.75 min^−1^), the decrease in network activation gradually disappears (compare **Figure S5C – D** to **Figure S5B**). Note that although the maximum in network activation does not change, its shifts to the right, requiring more ligands to reach maximal network activation (compare dashed line in **Figures S5B – D**).

Given the mechanistic detail of the model, we can explain the decrease in network activation observed upon mono-stimulation of NKG2D (**Figure 4**; orange star). As the ligand concentration increases, the velocity of phospho-NKG2D activation increases proportionally due to Michaelis-Menten kinetics. Based on the numerical value of *k*_*cat*_*_pNKG2D_pSHP*, the rate of dephosphorylation of phospho-NKG2D will be slow (or fast) if the parameter is small (or large). If the rate of dephosphorylation is slow, then the concentration of phospho-NKG2D will increase rapidly as the ligand concentration increases, and since there is a first-order degradation reaction for the phospho-species in our model, phospho-NKG2D will degrade proportionally to its concentration. Thus, when phospho-NKG2D activation is too fast, it will also decay rapidly, which will impede downstream signaling. This is why when the ligand concentration becomes too large, it has a suboptimal effect on network activation. Contrastingly, if the rate of dephosphorylation is fast, then the concentration of phospho-NKG2D will increase very slowly, which in turn will delay phospho-NKG2D decay and enable downstream signaling to continue.

Overall, these results suggest the magnitude in network activation is dependent on the receptor concentrations. Moreover, the ligand concentration needed to attain the maximal response is sensitive to the rate of phospho-receptor dephosphorylation, where the *faster* the phospho-receptor is dephosphorylated, the *greater* the ligand concentration needs to be to reach maximal activation. Taken together, the results presented here underscore the model’s utility in explaining and characterizing the system’s response to variations in input intensity.

### 3.4 Co-stimulation of CD16 and NKG2D potently activates the network

The impact on network activation induced by the co-stimulation of NK cell receptors has not been completely characterized. This knowledge gap obscures our understanding of how signals from multiple pathways are integrated and influence the downstream species. Thus, we simulated the model to better understand how co-stimulation affects network activation in a dose-dependent manner. Similar to the previous section, we varied the ligand concentration from 6.67 × 10^−5^ *μ*M up to 66.7 *μ*M and simulated the model for 60 minutes to observe how the magnitude of network activation changes.

Interestingly, the co-stimulation of CD16 and NKG2D (**Figure 5A**, red line) achieves greater network activation when compared to other combinations at equal ligand concentrations. With the exception of 2B4 and NKG2D costimulation (**Figure 5A**, magenta line), all combinations attain maximal network activation (116 *μ*M × min). In addition, we wanted to determine how much ligand is required to reach half-maximal network activation (**Figure 5A**, dashed line), akin to half-maximal effective concentration (EC_50_). We applied the model to predict the ligand concentration needed to reach this level of network activation for each combination (**Figure 5B**). The co-stimulation of CD16 and NKG2D (**Figure 5B**, red bar) required 26%, 27% and 51% fewer ligands on average compared to the ligand concentration needed for half-maximal activation with co-stimulation of all pathways (**Figure 5B**, black bar), 2B4 and NKG2D (**Figure 5B**, magenta bar), and CD16 and 2B4 (**Figure 5B**, green bar), respectively. At first glance, one may think that stimulating all pathways together would require the lowest ligand concentration to reach half-maximal network activation. However, since phospho-2B4 can activate phosphatases in addition to the kinase pSFK (**Figure 1**), co-stimulation of all three pathways is less effective than the co-stimulation of CD16 and NKG2D due to more phosphatase activation. In summary, we found that the co-stimulation CD16 and NKG2D *in silico* is more potent in activating the stimulatory network than all other combinations.

**Fig. 5.**
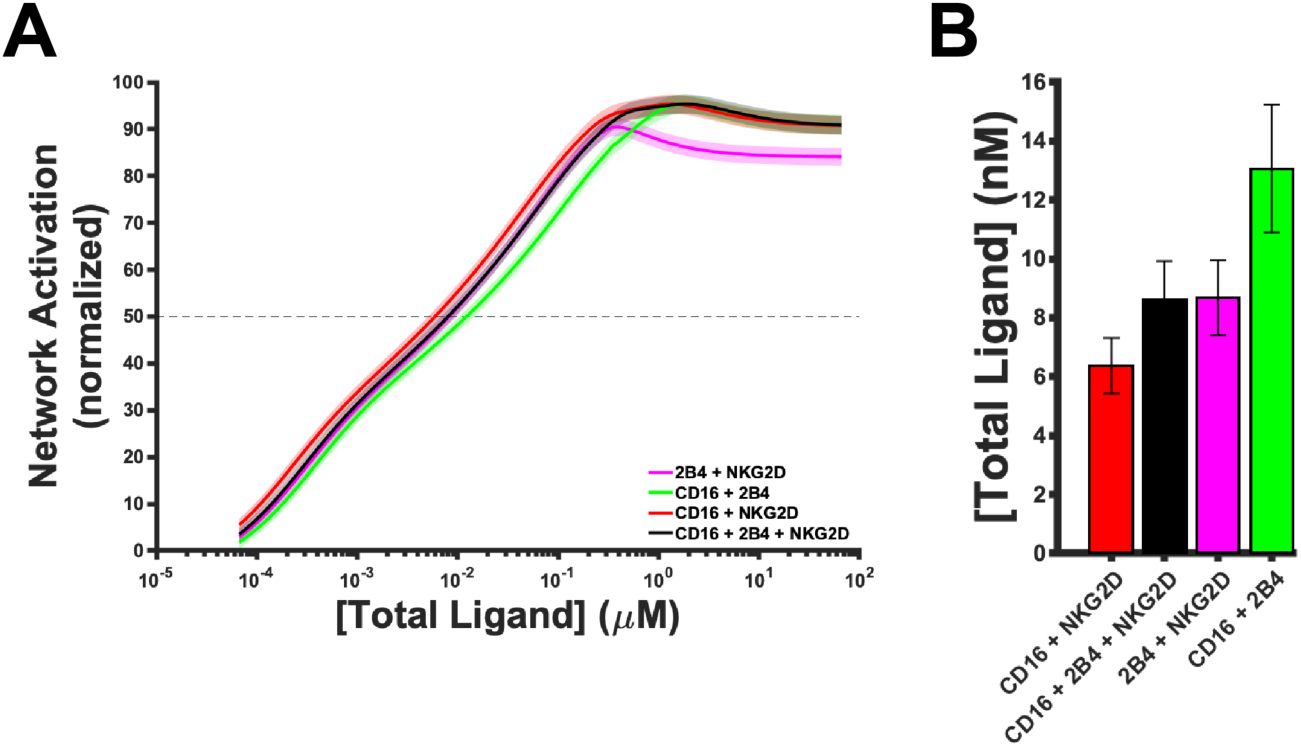
Receptor co-stimulation. Each receptor combination is stimulated with varying concentrations of ligands. (**A**) The co-stimulation of 2B4 and NKG2D (magenta), CD16 and 2B4 (green), CD16 and NKG2D (red) as well as the stimulation of all three receptors (black) are shown. The solid line represents the mean value from the 1,000 parameter estimates and the shaded area is one standard deviation. Network activation was scaled onto a range of [0,100] by normalizing the network activation by the maximum value across all three pathways. (**B**) The ligand concentration required to reach half-maximal activation (dashed line in panel **A**). Bars represent the mean model prediction from the 1,000 parameter estimates and the error bars represent one standard deviation.

### 3.5 In silico perturbations highlight the role of phospho-receptors and phosphatases in enhancing network activation

Mathematical models are instrumental in studying the trajectory of dynamical systems, especially when perturbations are considered. For example, model parameters can be varied to understand the system’s response to specific alterations. Therefore, we simulated the following perturbations to understand which changes augment network activation: (1) decreasing the rate of pSFK deactivation, (2) inhibiting pSHP activity, (3) increasing receptor-ligand affinity and (4) decreasing the decay rate of the phospho-receptors.

The first perturbation is inspired by experimental results (41,42) where a mutation of the activation-loop tyrosine (Y394) of lymphocyte-specific protein kinase (LCK), a member of the Src family kinase (SFK), disables the kinase from being inactivated in the context of T cell receptor signaling. This mutation evidently enhances T cell activation. Likewise, inhibiting the rate at which pSFK is deactivated (i.e., decreasing *k*_*cat*_*_pSFK_pSHP*) in the model may increase NK cell network activation by the same reasoning. It is known that phosphatases play an integral role in inhibiting NK cell activation (31,43–45) by dephosphorylating the downstream species. Therefore, inhibiting phosphatase activity (i.e., decreasing *k*_cat__pX_pSHP, where pX is a substrate for pSHP) is another mechanism that can increase network activation. Moreover, increasing the binding affinity between the ligand and the receptor should increase the velocity at which the receptor-ligand complex is formed, and thereby allow signaling to proceed more rapidly and possibly increase the magnitude of network activation. We simulated this effect by decreasing the *k*_*off*_ constant between the receptor and the ligand in the model. Finally, we decreased the decay rate of the phospho-receptors (i.e., *k*_*deg*_), as another means of modulating the network activation. This is inspired by Spran *et al.* (46), where they inhibited the shedding of CD16 receptors by introducing a point-wise amino acid mutation (S197P) that renders the receptor insusceptible to ADAM17-mediated cleavage. This engineered receptor induced more perforin degranulation upon stimulation, which is a downstream response of network activation. Although each of the perturbations should increase network activation in their own right, it is not obvious which perturbation (and to which extent) is the best approach. Thus, we simulated each case to determine which method is optimal for augmenting the magnitude of network activation.

We varied the parameters regulating the four perturbations from their baseline values up to 10-fold. The model was simulated for 60 minutes using various ligand concentrations. As before, we used the last 1,000 iterations from the best parameter set to simulate the model. Moreover, we simulated each perturbation separately for each pathway in order to observe any differences (or similarities) in the effects each perturbation imposes on each pathway. The simulated results can be found in **Figure 6**, where the percent change in network activation from baseline via mono-stimulation of NKG2D (**Figure 6A – C**), CD16 (**Figure 6D –F**) and 2B4 (**Figure 6G – I**) is plotted as a function of the change in the strength of the perturbation. The circles, triangles, squares and diamonds in **Figure 6** correspond to decreasing pSFK deactivation, inhibiting pSHP activity, increasing ligand affinity and inhibiting phospho-receptor decay, respectively.

**Fig. 6.**
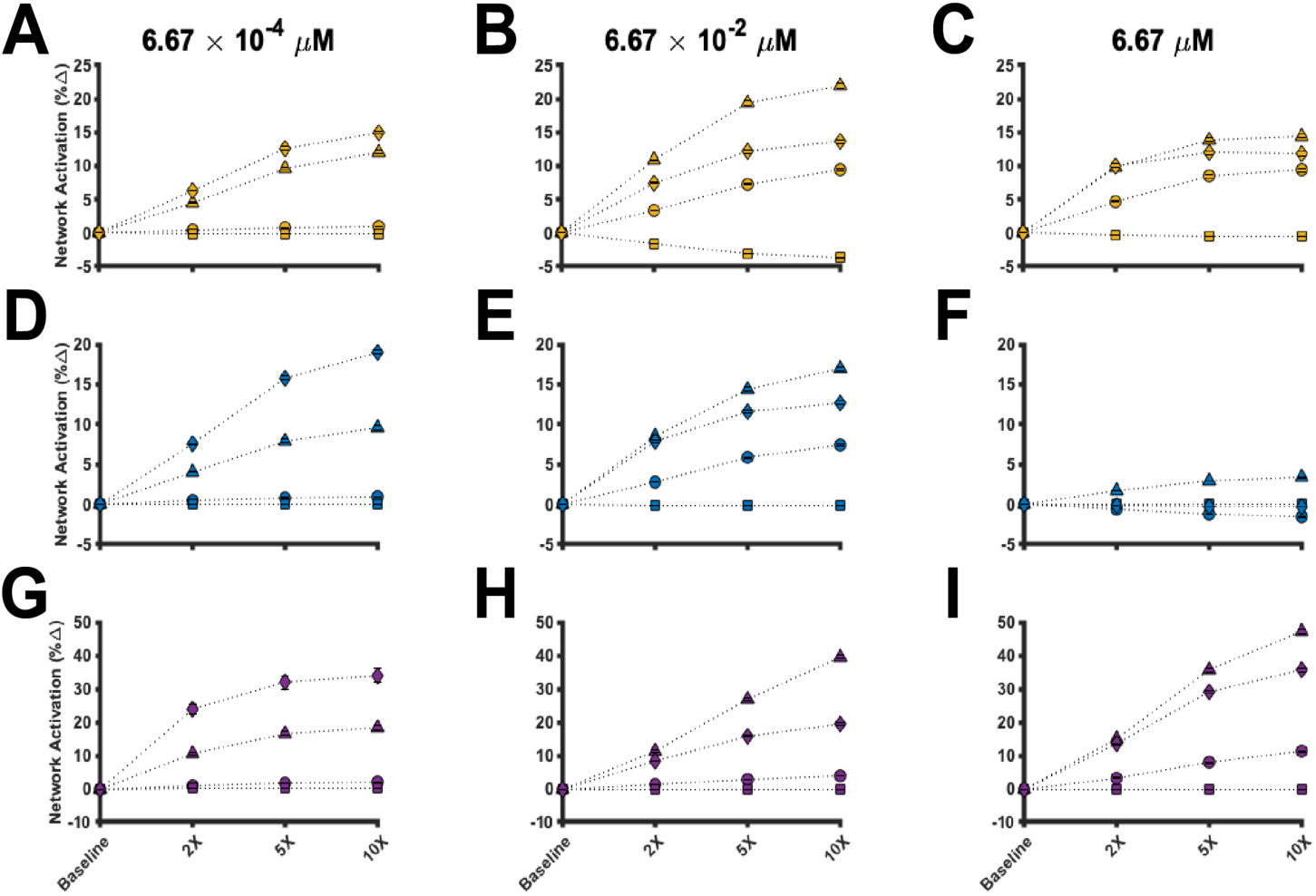
Perturbations to the stimulatory network. The percent change of the magnitude of network activation from baseline for stimulation of NKG2D (**A – C**), CD16 (**D – F**) and 2B4 (**G – I**) is shown. The perturbations were simulated using a ligand concentration of 6.67 × 10^−4^ *μ*M (left column), 6.67 × 10^−2^ *μ*M (middle column) and 6.67 *μ*M (right column). Circles: decreasing pSFK deactivation rate. Triangles: decreasing pSHP activity. Squares: increasing receptor-ligand affinity. Diamonds: decreasing phospho-receptor decay rate. Marker: mean value from 1,000 parameter estimates. Error bars: one standard deviation.

The model provides detailed insight into the effects of perturbing the signaling network. Firstly, when the ligand concentration is low (**Figure 6A, D** and **G**), the percent change in network activation is more sensitive to phosphoreceptor decay (diamonds). In contrast, when the ligand concentration is moderate (**Figure 6B, E** and **H**) to high (**Figure 6C, F** and **I**), the percent change in network activation is influenced more by phosphatase activity (triangles). These results hold true for all pathways, suggesting the perturbations qualitatively impact the pathways in a similar manner. Interestingly, when considering the NKG2D and CD16 pathways, the relative effect of the perturbations decreases as the concentration of the input increases (compare **Figure 6C** to **Figure 6B** and compare **Figure 6F** to **Figure 6E**). Surprisingly, the relative impact of decreasing pSFK deactivation (circles) and increasing ligand affinity (squares) on network activation is almost negligible. In some cases, increasing ligand affinity can even decrease network activation.

These data suggest network activation is tightly controlled by the phospho-receptors and the phosphatases. Based on the model predictions, when input levels are low, it is more important to engineer receptors that are resistant to proteolytic cleavage, as this enables the activated receptor to induce continued intracellular signaling. Alternatively, when the input to the system is plentiful, inhibiting phospho-receptor decay is not influential since the large concentration of input can enable continued intracellular signaling. In this scenario, it is more influential to inhibit phosphatase activity, which allows the phospho-species to remain activated, thereby increasing the magnitude of species activation. In summary, the model predicts that the phospho-receptors and phosphatases strongly regulate the magnitude of network activation, and that the optimal strategy for enhancing network activation is dependent on the level of stimulation. Thus, our simulations provide quantitative insight into mechanisms that can augment the activation of the stimulatory network in NK cells.

## DISCUSSION

In the present study, we constructed a mathematical model of a subset of the signaling pathways that mediate NK cell stimulation. We interrogated the model to understand (1) how the stimulatory network is influenced by the different pathways, (2) which signaling species and parameters directly influence the magnitude of network activation, (3) which combination of receptors are more potent in activating the stimulatory network and (4) how the network can be perturbed to enhance activation.

Our baseline model predictions demonstrate network activation is sensitive to both the receptor concentrations as well as the rate of receptor deactivation. Specifically, the receptor concentrations influence the magnitude of activation whereas the rate of receptor deactivation influences the ligand concentration needed to attain a certain level of network activation. From our perturbation studies, we observed that the phospho-receptors and the phosphatases control the system’s response to NK cell receptor stimulation. Inhibiting phospho-receptor decay is particularly important for enhancing network activation when the input to the system is scarce. Alternatively, when the input is abundant, it is more important to inhibit phosphatase activity. In the case where NK cells are directed to recognize specific tumor-associated antigen via engineered receptors such as chimeric antigen receptors (CARs), our simulations suggest engineering receptors to be resistant to proteolytic cleavage, as the antigen may not be abundantly expressed on the tumor cell surface. In monoclonal antibody therapies, which can expose NK cells to a large concentration of input, the model simulations indicate that pre-incubation of NK cells with pan-SHP inhibitors may unbridle NK cell activation and allow for a strong response. Thus, our computational modeling of NK cell stimulation is highly valuable and particularly useful. Besides the large amount of time and resources needed to complete such studies via experimentation alone, many of the nonlinear properties embedded in the signaling network would be difficult to capture and effectively characterize without knowledge of the parameters regulating the system. Instead, when there is a healthy union between data and modeling, our understanding of biology benefits the most.

Our modeling results provide a robust quantitative framework to study the effects of co-stimulation of NK cell receptors. These predictions are relevant for developing immunotherapeutic strategies. Researchers in recent years have designed CARs for NK cells that include intracellular signaling domains of CD16, 2B4 and NKG2D for anti-tumor therapy (47,48). Those studies found that CARs comprised of CD16, 2B4 and NKG2D signaling domains together outperformed activation induced by the individual receptors. In addition, CAR-NK cell immunotherapies (49,50) that include intracellular domains from both CD16 and NKG2D are shown to be effective in eliminating tumors in pre-clinical studies. Through continued success in the pre-clinical stage, a few CAR-NK cell immunotherapies have entered clinical trials as potential therapeutics for cancer patients (51,52). Excitingly, our model predicts that co-stimulation of CD16 and NKG2D activate the network strongly both individually and collectively. We infer from our results that CARs that express the signaling domains of CD16 (CD3ζ) and NKG2D (DAP10) may promote strong activation of the signaling network. Although the model presented here was not trained on data from CARs, our results are in accord with those found by researchers in immunotherapy. This demonstrates the model’s utility in predicting which strategies can improve NK cell activation. Importantly, the modeling predictions go beyond observations from published experimental studies by providing detailed predictions about the magnitude of activation across a range of ligand concentrations and insight as to why certain combinations work better than others.

We acknowledge some limitations that may affect the model predictions. Firstly, our model includes three important stimulatory receptors; however, several others could have been considered as well. Additionally, although multiple sites of phosphorylation and dephosphorylation can exist for each species, we have not included this level of detail in the model. This would increase the specificity of our model, but it would be at the expense of model simplicity. Since we are interested in understanding and comparing the dynamics between multiple pathways, we sought to retain a simplified model in order to effectively compare the pathways. In the future, researchers can adopt and improve the current model by considering site specific reactions and their importance in particular aspects of NK cell stimulation. Finally, although the initial concentrations of the signaling species were derived from literature (20–23), we expect that these values may differ based on the specific NK cell line or the donor for primary NK cells. Future research can address these limitations, building upon the work presented here. In addition, questions within tumor immunology, in particular tumor and NK cell dynamics, can be studied by integrating the present signaling model with a cell-based model. Mahasa and coworkers provide an example of such a model that incorporates intracellular and intercellular dynamics (11).

Despite these limitations, our mathematical model is relevant in understanding NK cell signaling and how the stimulatory network can be enhanced. The results presented here lend support for multiple strategies to induce optimal cell activation, including (1) the co-stimulation of specific receptor combinations, which is relevant to the design of engineered receptors (e.g., CARs), (2) modifying such engineered receptors to be degradation-resistant to promote continued signaling and (3) inhibiting SHP activity, when input levels are sufficiently large, to disinhibit the activation of the signaling species that contribute to cell activation. In conclusion, our work provides strategies and insight into engineering NK cells for enhanced activation.

## ACKNOWLEDGEMENTS

We immensely appreciate the Finley research group for critical evaluation of this manuscript as well as improvements to the model code. This work was supported a Viterbi/Graduate School Merit Fellowship (to S.Z.M).

## SUPPORTING INFORMATION

**File S1.** Computational model files (.m files with model code and .mat files with parameter values); provided as .zip file.

**File S2.** List of model species, reactions and parameters; provided as .xlsx file.

**File S3.** Experimental data and description of data processing required for model calibration and validation; provided as .xlsx file.

**File S4.** Experimental data and description of data processing required for model calibration and validation; provided as .xlsx file.

## Notes

#### Summary of Updates

- Performed new parameter estimation using a Bayesian approach - Performed new calculations to characterize the extent of network activation - Added results from simulating perturbations to the signaling network - New figures added (Fig. 2, 3, 4, 5, and 6) - Added supplemental file with all data used for model calibration and validation

