## Supplementary material for "Enhancing network activation in Natural Killer cells: Predictions from *in silico* modeling": File S1: Driver.m

```
%%%%%%%%%%%%%%%
% Driver file %
%%%%%%%%%%%%%%%

close all;
clear;
clc;

%% Load Inputs

% Parameters
load('parameters.mat');

% Initial Concentrations
load('CD16.mat');
load('TwoB4.mat');
load('NKG2D.mat');

% Set ODE options
options = odeset('RelTol',1E-09,...
                    'AbsTol',1E-12,...
                    'NonNegative',1:36);


% Initialize Time Step
tspan = linspace(0,60,601)'; % 60 minutes

%% Simulate Model

parfor i = 1:size(parameters,2)
    
    [~, spec_conc_CD16(:,:,i)] = ode15s(@NK_CD16_ODE,tspan,y0_CD16,options,parameters(:,i));
    [~, spec_conc_2B4(:,:,i)] = ode15s(@NK_2B4_ODE,tspan,y0_2B4,options,parameters(:,i));
    [~, spec_conc_NKG2D(:,:,i)] = ode15s(@NK_NKG2D_ODE,tspan,y0_NKG2D,options,parameters(:,i));
    
end


    
```
