## Supplementary material for "Enhancing network activation in Natural Killer cells: Predictions from *in silico* modeling": File S1: NK_2B4_ODE.m

```
function dxdt = NK_Master_2B4_ODE(t,x,theta)

%% Define x

    CD16 = x(1); 
    LCD16 = x(2);
    CD16_LCD16 = x(3);
    pCD16_LCD16 = x(4);
    
    TwoB4 = x(5);
    L2B4 = x(6);
    TwoB4_L2B4 = x(7);
    p2B4_L2B4 = x(8);
    SAP_p2B4_L2B4 = x(9);
    
    NKG2D = x(10);
    LNKG2D = x(11);
    NKG2D_LNKG2D = x(12);
    pNKG2D_LNKG2D = x(13);
    
    SAP = x(14);
    
    SFK = x(15);
    pSFK = x(16);
    
    ZAP70 = x(17);
    pZAP70 = x(18);
    
    LAT = x(19);
    pLAT = x(20);
    
    PLCg = x(21);
    pPLCg = x(22);
    
    Akt = x(23);
    pAkt = x(24);
    
    SLP76 = x(25);
    pY113SLP76 = x(26);
    pY128SLP76 = x(27);
    ppSLP76 = x(28);
    
    Vav = x(29);
    pVav = x(30);
    
    Erk = x(31);
    pErk = x(32);
    
    SHP = x(33);
    pSHP = x(34);
    
    SHIP = x(35);
    pSHIP = x(36);
    
%% Define theta

%% Receptor Ligand Interactions

    kon_CD16_LCD16 = theta(1);
    koff_CD16_LCD16 = theta(2);
    
    kon_2B4_L2B4 = theta(3);
    koff_2B4_L2B4 = theta(4);
    
    kon_NKG2D_LNKG2D = theta(5);
    koff_NKG2D_LNKG2D = theta(6);
    
%% Receptor activation/deactivation
    
    kcat_CD3z_SFK = theta(7); 
    Km_CD3z_SFK = theta(8);
    
    kcat_pCD3z_pSHP = theta(9);
    Km_pCD3z_pSHP = theta(10);
    
    kcat_2B4_SFK = theta(11); 
    Km_2B4_SFK = theta(12);
    
    kcat_p2B4_pSHP = theta(13); 
    Km_p2B4_pSHP = theta(14);
    
    kcat_NKG2D_SFK = theta(15); 
    Km_NKG2D_SFK = theta(16);
    
    kcat_pNKG2D_pSHP = theta(17);
    Km_pNKG2D_pSHP = theta(18);
    
%% SAP Binding to p2B4
    
    kon_p2B4_SAP = theta(19);
    koff_p2B4_SAP = theta(20);
    
%% SFK Activation/Deactivation
    
    kcat_SFK_pCD3z = theta(21);
    Km_SFK_pCD3z = theta(22);
    
    kcat_SFK_SAP = theta(23); 
    Km_SFK_SAP = theta(24);
    
    kcat_SFK_pNKG2D = theta(25);
    Km_SFK_pNKG2D = theta(26);
    
    kcat_pSFK_pSHP = theta(27);
    Km_pSFK_pSHP = theta(28);
    
%% ZAP70 Activation/Deactivation

    kcat_ZAP70_pSFK = theta(29);
    Km_ZAP70_pSFK = theta(30);
    
    kcat_pZAP70_pSHP = theta(31);
    Km_pZAP70_pSHP = theta(32);
    
%% LAT Activation/Deactivation

    kcat_LAT_pSFK = theta(33);
    Km_LAT_pSFK = theta(34);

    kcat_LAT_pZAP70 = theta(35);
    Km_LAT_pZAP70 = theta(36);
    
    kcat_pLAT_pSHP = theta(37);
    Km_pLAT_pSHP = theta(38);
    
%% Akt Activation/Deactivation

    kcat_uAkt_pSFK = theta(39);
    Km_uAkt_pSFK = theta(40);
    
    kcat_pAkt_pSHIP = theta(41);
    Km_pAkt_pSHIP = theta(42);
    
%% PLCg Activation/Deactivation

    kcat_PLCg_pSFK = theta(43);
    Km_PLCg_pSFK = theta(44);
    
    kcat_pPLCg_pSHP = theta(45);
    Km_pPLCg_pSHP = theta(46);
    
%% Vav Activation/Deactivation

    kcat_Vav_pSFK = theta(47);
    Km_Vav_pSFK = theta(48);
    
    kcat_pVav_pSHP = theta(49);
    Km_pVav_pSHP = theta(50);
    
%% SLP76 Activation/Deactivation

    kcat_p113SLP76_pSFK = theta(51);
    Km_p113SLP76_pSFK = theta(52);
    
    kcat_p128SLP76_pSFK = theta(53);
    Km_p128SLP76_pSFK = theta(54);
    
    kcat_p113SLP76_p128_pSFK = theta(55);
    Km_p113SLP76_p128_pSFK = theta(56);

    kcat_p128SLP76_p113_pSFK = theta(57);
    Km_p128SLP76_p113_pSFK = theta(58);
    
    kcat_p113SLP76_p113_pSHP = theta(59);
    Km_p113SLP76_p113_pSHP = theta(60);
    
    kcat_p128SLP76_p128_pSHP = theta(61);
    Km_p128SLP76_p128_pSHP = theta(62);
    
    kcat_p113SLP76_fp_pSHP = theta(63);
    Km_p113SLP76_fp_pSHP = theta(64);

    kcat_p128SLP76_fp_pSHP = theta(65);
    Km_p128SLP76_fp_pSHP = theta(66);
    
%% Erk Activation/Deactivation

    kcat_uErk_pErk = theta(67);
    Km_uErk_pErk = theta(68);

    kcat_pErk_pSHP = theta(69);
    Km_pErk_pSHP = theta(70);
    
%% SHP Activation/Deactivation

    kcat_SHP_pSFK = theta(71);
    Km_SHP_pSFK = theta(72);

    kcat_SHP_p2B4 = theta(73); 
    Km_SHP_p2B4 = theta(74);
    
    kcat_pSHP_pSHP = theta(75);
    Km_pSHP_pSHP = theta(76);
    
%% SHIP Activation/Deactivation
    
    kcat_SHIP_pSFK = theta(77);
    Km_SHIP_pSFK = theta(78);
    
    kcat_SHIP_p2B4 = theta(79); 
    Km_SHIP_p2B4 = theta(80);

    kcat_pSHIP_pSHP = theta(81);
    Km_pSHIP_pSHP = theta(82);
    
%% Decay

    kdeg = theta(83);
        
%% Functions

    MM_SFK_pNKG2D = ((kcat_SFK_pNKG2D)*(pNKG2D_LNKG2D)*SFK)/(Km_SFK_pNKG2D + SFK);

    MM_pNKG2D_pSHP = ((kcat_pNKG2D_pSHP)*pSHP*pNKG2D_LNKG2D)/(Km_pNKG2D_pSHP + pNKG2D_LNKG2D);

    MM_NKG2D_SFK = ((kcat_NKG2D_SFK)*(NKG2D_LNKG2D)*(SFK))/(Km_NKG2D_SFK + NKG2D_LNKG2D);

    MM_2B4_SFK = ((kcat_2B4_SFK)*(TwoB4_L2B4)*(SFK))/(Km_2B4_SFK + TwoB4_L2B4);

    MM_p2B4_pSHP = ((kcat_p2B4_pSHP)*pSHP*p2B4_L2B4)/(Km_p2B4_pSHP + p2B4_L2B4);
    
    MM_SFK_SAP = ((kcat_SFK_SAP)*(SAP_p2B4_L2B4)*SFK)/(Km_SFK_SAP + SAP_p2B4_L2B4);
    
    MM_SHP_p2B4 = ((kcat_SHP_p2B4)*(p2B4_L2B4)*SHP)/(Km_SHP_p2B4 + p2B4_L2B4);
    
    MM_SHIP_p2B4 = ((kcat_SHIP_p2B4)*(p2B4_L2B4)*SHIP)/(Km_SHIP_p2B4 + p2B4_L2B4);
    
    MM_CD3z_SFK = ((kcat_CD3z_SFK)*(CD16_LCD16)*(SFK))/(Km_CD3z_SFK + CD16_LCD16);

%     MM_ZAP70_pSFK = ((kcat_ZAP70_pSFK)*(pSFK)*(ZAP70))/(Km_ZAP70_pSFK + ZAP70);

%     MM_LAT_pSFK = ((kcat_LAT_pSFK)*pSFK*LAT)/(Km_LAT_pSFK + LAT);

%     MM_LAT_pZAP70 = ((kcat_LAT_pZAP70)*pSFK*LAT)/(Km_LAT_pZAP70 + LAT);

    MM_p113SLP76_pSFK = ((kcat_p113SLP76_pSFK)*pSFK*SLP76)/(Km_p113SLP76_pSFK + SLP76);

%     MM_p113SLP76_p128_pSFK = ((kcat_p113SLP76_p128_pSFK)*pSFK*pY128SLP76)/(Km_p113SLP76_p128_pSFK + pY128SLP76);

%     MM_p128SLP76_pSFK = ((kcat_p128SLP76_pSFK)*pSFK*SLP76)/(Km_p128SLP76_pSFK + SLP76);

%     MM_p128SLP76_p113_pSFK = ((kcat_p128SLP76_p113_pSFK)*pSFK*pY113SLP76)/(Km_p128SLP76_p113_pSFK + pY113SLP76);

    MM_Vav_pSFK = ((kcat_Vav_pSFK)*pSFK*Vav)/(Km_Vav_pSFK + Vav);

    MM_SHP_pSFK = ((kcat_SHP_pSFK)*pSFK*SHP)/(Km_SHP_pSFK + SHP);

    MM_SHIP_pSFK = ((kcat_SHIP_pSFK)*pSFK*SHIP)/(Km_SHIP_pSFK + SHIP);

    MM_pSFK_pSHP = ((kcat_pSFK_pSHP)*pSHP*pSFK)/(Km_pSFK_pSHP + pSFK);

    MM_pZAP70_pSHP = ((kcat_pZAP70_pSHP)*pSHP*pZAP70)/(Km_pZAP70_pSHP + pZAP70);

    MM_pCD3z_pSHP = ((kcat_pCD3z_pSHP)*pSHP*pCD16_LCD16)/(Km_pCD3z_pSHP + pCD16_LCD16);

    MM_pLAT_pSHP = ((kcat_pLAT_pSHP)*pSHP*pLAT)/(Km_pLAT_pSHP + pLAT);

    MM_pVav_pSHP = ((kcat_pVav_pSHP)*pSHP*pVav)/(Km_pVav_pSHP + pVav);

    MM_pPLCg_pSHP = ((kcat_pPLCg_pSHP)*pSHP*pPLCg)/(Km_pPLCg_pSHP + pPLCg);

    MM_p113SLP76_p113_pSHP = ((kcat_p113SLP76_p113_pSHP)*pSHP*pY113SLP76)/(Km_p113SLP76_p113_pSHP + pY113SLP76);

    MM_p113SLP76_fp_pSHP = ((kcat_p113SLP76_fp_pSHP)*pSHP*ppSLP76)/(Km_p113SLP76_fp_pSHP + ppSLP76);

    MM_p128SLP76_p128_pSHP = ((kcat_p128SLP76_p128_pSHP)*pSHP*pY128SLP76)/(Km_p128SLP76_p128_pSHP + pY128SLP76);

    MM_p128SLP76_fp_pSHP = ((kcat_p128SLP76_fp_pSHP)*pSHP*ppSLP76)/(Km_p128SLP76_fp_pSHP + ppSLP76);

    MM_PLCg_pSFK = ((kcat_PLCg_pSFK)*(pSFK)*PLCg)/(Km_PLCg_pSFK + PLCg);

    MM_uErk_pErk = ((kcat_uErk_pErk)*(pY113SLP76 + pVav)*Erk)/(Km_uErk_pErk + Erk);

    MM_pErk_pSHP = ((kcat_pErk_pSHP)*pSHP*pErk)/(Km_pErk_pSHP + pErk);

%     MM_uAkt_pSFK = ((kcat_uAkt_pSFK)*(pSFK)*Akt)/(Km_uAkt_pSFK + Akt);

    MM_pAkt_pSHIP = ((kcat_pAkt_pSHIP)*(pSHIP)*pAkt)/(Km_pAkt_pSHIP + pAkt);

    MM_pSHP_pSHP = ((kcat_pSHP_pSHP)*(pSHP)*pSHP)/(Km_pSHP_pSHP + pSHP);

    MM_pSHIP_pSHP = ((kcat_pSHIP_pSHP)*(pSHP)*pSHIP)/(Km_pSHIP_pSHP + pSHIP);

    MM_SFK_pCD3z = ((kcat_SFK_pCD3z)*(pCD16_LCD16)*SFK)/(Km_SFK_pCD3z + SFK);


%% x Derivatives

%% CD16

    % CD16
    dxdt(1,1) = (koff_CD16_LCD16*CD16_LCD16) - (kon_CD16_LCD16*CD16*LCD16);
    
    % Ligand CD16
    dxdt(2,1) = (koff_CD16_LCD16*CD16_LCD16) - (kon_CD16_LCD16*CD16*LCD16);    
    
    % CD16:Ligand CD16
    dxdt(3,1) = (kon_CD16_LCD16*CD16*LCD16) + (MM_pCD3z_pSHP) - (koff_CD16_LCD16*CD16_LCD16) - (MM_CD3z_SFK);

    % pCD16:Ligand CD16
    dxdt(4,1) = MM_CD3z_SFK - MM_pCD3z_pSHP - kdeg*pCD16_LCD16;
    
%% 2B4
    
    % 2B4
    dxdt(5,1) = (koff_2B4_L2B4*TwoB4_L2B4) - (kon_2B4_L2B4*TwoB4*L2B4);
    
    % Ligand
    dxdt(6,1) = (koff_2B4_L2B4*TwoB4_L2B4) - (kon_2B4_L2B4*TwoB4*L2B4);    
    
    % 2B4:Ligand
    dxdt(7,1) = (kon_2B4_L2B4*TwoB4*L2B4) + (MM_p2B4_pSHP) - (koff_2B4_L2B4*TwoB4_L2B4) - (MM_2B4_SFK);

    % p2B4:Ligand
    dxdt(8,1) = MM_2B4_SFK - MM_p2B4_pSHP - (kon_p2B4_SAP*p2B4_L2B4*SAP) + (koff_p2B4_SAP*SAP_p2B4_L2B4) - kdeg*p2B4_L2B4;
    
    % SAP:p2B4:Ligand
    dxdt(9,1) = (kon_p2B4_SAP*p2B4_L2B4*SAP) - (koff_p2B4_SAP*SAP_p2B4_L2B4) - kdeg*SAP_p2B4_L2B4;
    
%% NKG2D
    
    % NKG2D
    dxdt(10,1) = (koff_NKG2D_LNKG2D*NKG2D_LNKG2D) - (kon_NKG2D_LNKG2D*NKG2D*LNKG2D);
    
    % Ligand
    dxdt(11,1) = (koff_NKG2D_LNKG2D*NKG2D_LNKG2D) - (kon_NKG2D_LNKG2D*NKG2D*LNKG2D);    
    
    % NKG2D:Ligand
    dxdt(12,1) = (kon_NKG2D_LNKG2D*NKG2D*LNKG2D) + (MM_pNKG2D_pSHP) - (koff_NKG2D_LNKG2D*NKG2D_LNKG2D) - (MM_NKG2D_SFK);

    % pNKG2D:Ligand
    dxdt(13,1) = MM_NKG2D_SFK - MM_pNKG2D_pSHP - kdeg*pNKG2D_LNKG2D;
    
%% SAP
    
    % SAP
    dxdt(14,1) = (koff_p2B4_SAP*SAP_p2B4_L2B4) - (kon_p2B4_SAP*p2B4_L2B4*SAP);
    
%% SFK
    
    % SFK
    dxdt(15,1) = MM_pSFK_pSHP - MM_SFK_pCD3z - MM_SFK_SAP - MM_SFK_pNKG2D;
    
    % pSFK
    dxdt(16,1) = MM_SFK_pCD3z + MM_SFK_SAP + MM_SFK_pNKG2D - MM_pSFK_pSHP - pSFK*kdeg;
    
%% ZAP70
    
    % ZAP70
    dxdt(17,1) = MM_pZAP70_pSHP;
    
    % pZAP70
    dxdt(18,1) = - MM_pZAP70_pSHP - pZAP70*kdeg;
    
%% LAT
    
    % LAT
    dxdt(19,1) = MM_pLAT_pSHP;
    
    % pLAT
    dxdt(20,1) = - MM_pLAT_pSHP - pLAT*kdeg;
    
%% PLCg
    
    % PLCg
    dxdt(21,1) = MM_pPLCg_pSHP - MM_PLCg_pSFK;
    
    % pPLCg
    dxdt(22,1) = MM_PLCg_pSFK - MM_pPLCg_pSHP - pPLCg*kdeg;
    
%% Akt
    
    % Akt
    dxdt(23,1) = MM_pAkt_pSHIP;
    
    % pAkt
    dxdt(24,1) = - MM_pAkt_pSHIP - pAkt*kdeg;
    
%% SLP76
    
    % SLP76
    dxdt(25,1) = MM_p113SLP76_p113_pSHP + MM_p128SLP76_p128_pSHP - MM_p113SLP76_pSFK;
    
    % pY113 SLP76
    dxdt(26,1) = MM_p113SLP76_pSFK + MM_p128SLP76_fp_pSHP - MM_p113SLP76_p113_pSHP - pY113SLP76*kdeg;
    
    % pY128 SLP76
    dxdt(27,1) = MM_p113SLP76_fp_pSHP - MM_p128SLP76_p128_pSHP - pY128SLP76*kdeg;
    
    % ppSLP76
    dxdt(28,1) = - MM_p113SLP76_fp_pSHP - MM_p128SLP76_fp_pSHP - ppSLP76*kdeg;
    
%% Vav    

    % Vav
    dxdt(29,1) = MM_pVav_pSHP - MM_Vav_pSFK;
    
    % pVav
    dxdt(30,1) = MM_Vav_pSFK - MM_pVav_pSHP - pVav*kdeg;
    
%% Erk
    
    % Erk
    dxdt(31,1) = MM_pErk_pSHP - MM_uErk_pErk;
    
    % pErk
    dxdt(32,1) = MM_uErk_pErk - MM_pErk_pSHP - pErk*kdeg;
    
%% SHP
    
    % SHP
    dxdt(33,1) = MM_pSHP_pSHP - MM_SHP_pSFK - MM_SHP_p2B4;
    
    % pSHP
    dxdt(34,1) = MM_SHP_pSFK + MM_SHP_p2B4 - MM_pSHP_pSHP - pSHP*kdeg;
    
%% SHIP
    
    % SHIP
    dxdt(35,1) = MM_pSHIP_pSHP - MM_SHIP_p2B4 - MM_SHIP_pSFK;
    
    % pSHIP
    dxdt(36,1) = MM_SHIP_pSFK - MM_pSHIP_pSHP + MM_SHIP_p2B4 - pSHIP*kdeg;
    
%%
    
    return
```
