## Supplementary material for "Enhancing network activation in Natural Killer cells: Predictions from *in silico* modeling": File S4

Supplementary Figures

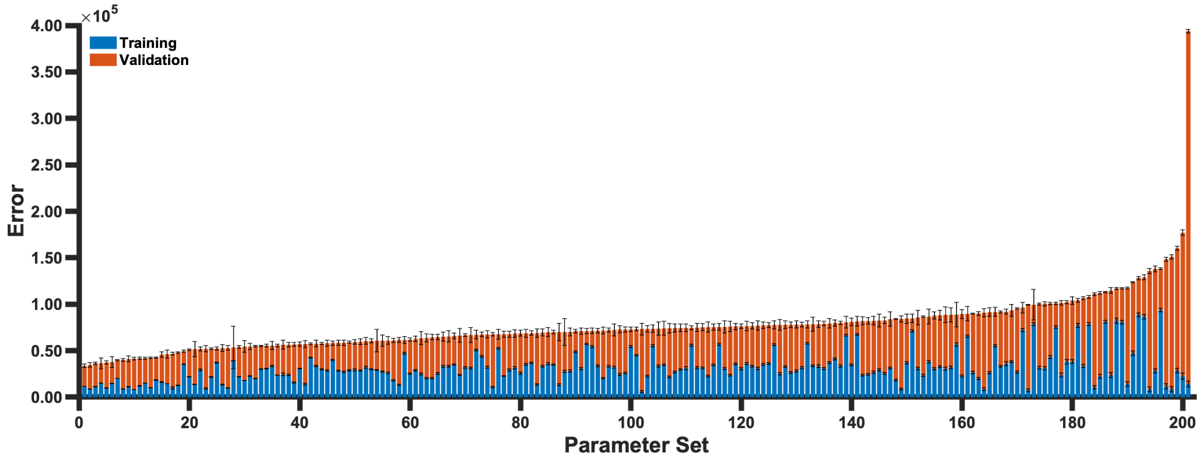


**Figure S1.** Total error from parameter estimation. The Metropolis-Hastings algorithm was simulated 200 times. For each run, we used the last 1,000 iterations to simulate the model and compare the model predictions to training (blue) and validation (red) data. Bars represent mean value and error bars represent one standard deviation. The total error was sorted in ascending order. The first 14 parameter sets were used to analyze the model predictions.

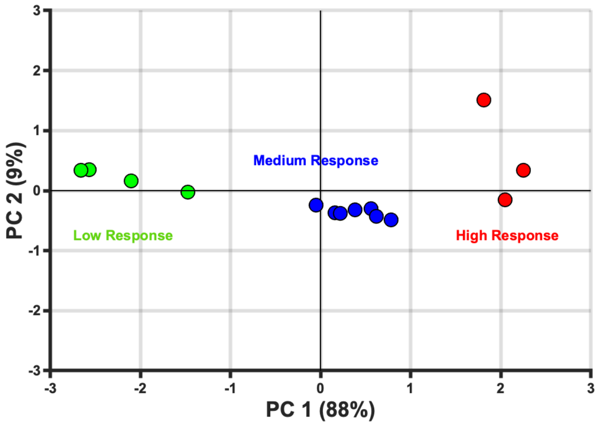


**Figure S2.** Cluster analysis of model predictions. We analyzed the model predictions using the 14 parameter sets. We clustered the model predictions using *kmeans* in MATLAB and found three clusters based on the magnitude of network activation (low, medium and high; green, blue and red, respectively). The model predictions were also analyzed using *pca* in MATLAB, where principal component 1 (PC 1) explained about 88% of the total variance in the model predictions. For both the *kmeans* and *pca* functions, the input was a 14 × 3 matrix corresponding to the mean model prediction from the 14 parameter sets using mono-stimulation of the three pathways.

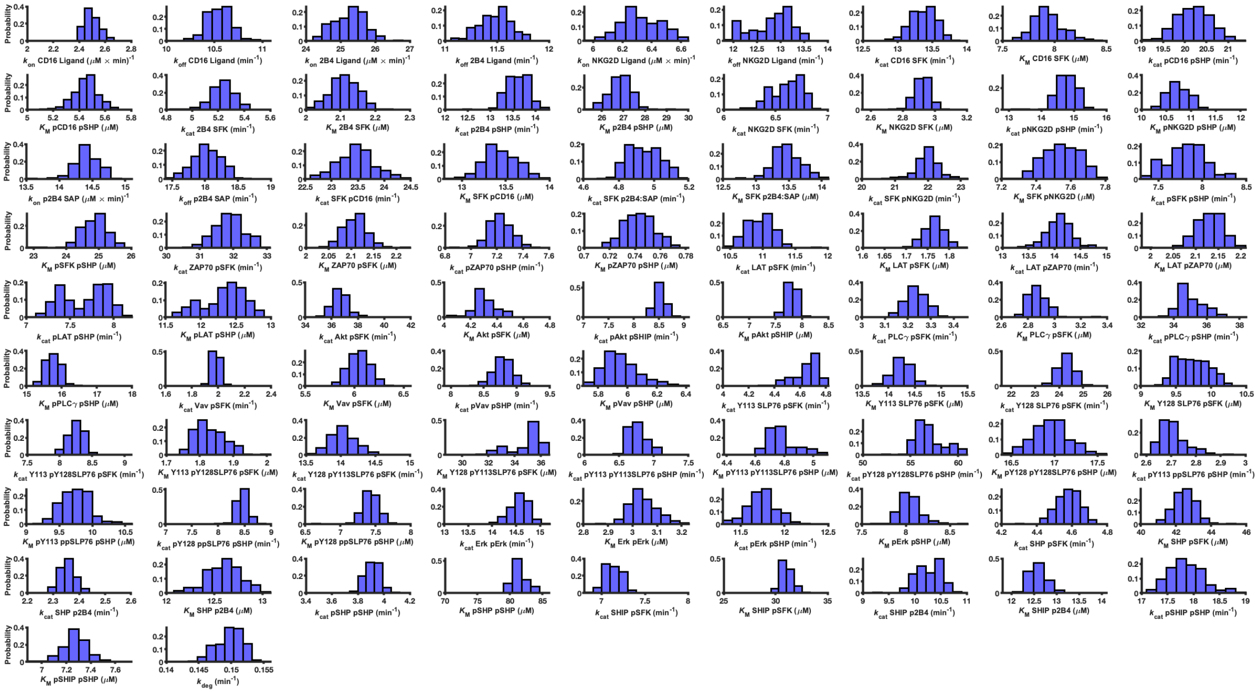

**Figure S3.** Parameter estimation using the best parameter set. The parameter distribution for each of the 83 estimated parameters are shown. We used the final 1,000 iterations from parameter estimation to plot the histograms for each parameter.

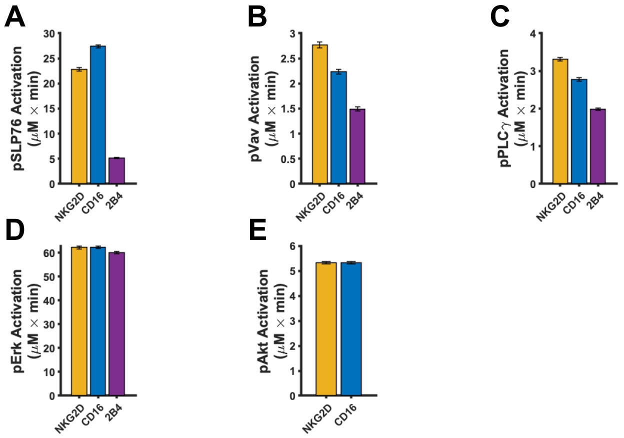


**Figure S4.** Magnitude of species activation. The magnitude of activation of the phospho-species (**A**) pSLP76, (**B**) pVav, (**C**) pPLC$\gamma$, (**D**) pErk and (**E**) pAkt is shown based on mono-stimulation of NKG2D (orange), CD16 (blue) and 2B4 (purple) using the 1,000 iterations from the best parameter set. The bar represents the mean value and the error bars represent one standard deviation.

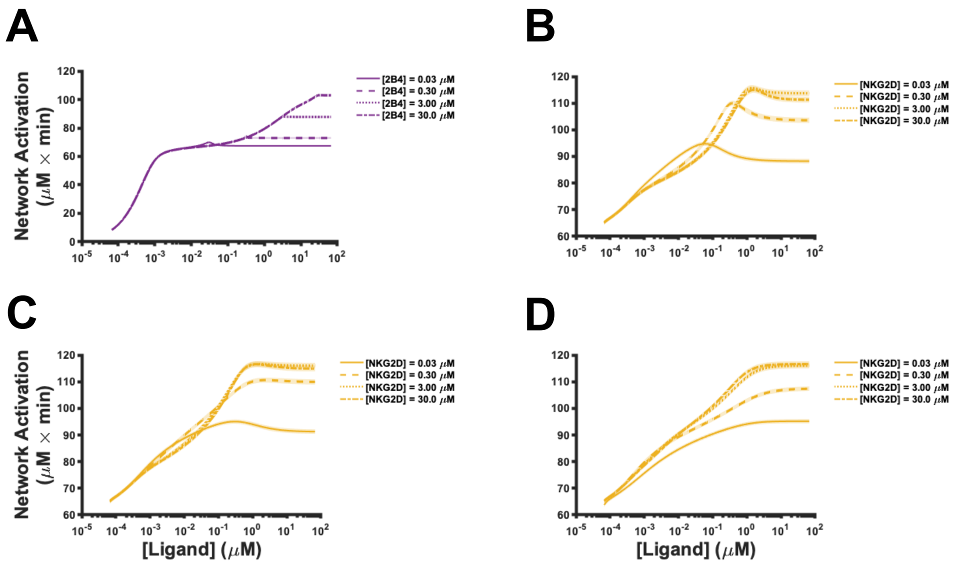


**Figure S5.** Effect of receptor concentrations on network activation. The (**A**) 2B4 and (**B – D**) NKG2D pathways were stimulated with ligands *in silico*. For all panels, the line represents the mean model prediction using the final 1,000 iterations from parameter estimation using the best set and the shaded area is one standard deviation. The solid, dash, dot and dash-dot lines correspond to 0.03, 0.3, 3 and 30 $\mu$M of the receptor, respectively.
